# A Driven Disordered Systems Approach to Biological Evolution in Changing Environments

**DOI:** 10.1101/2021.08.13.456229

**Authors:** Suman G Das, Joachim Krug, Muhittin Mungan

**Affiliations:** Institute for Biological Physics, University of Cologne, Zülpicher Straße 77, D-50937 Köln, Germany; Institut für Angewandte Mathematik, Universiät Bonn, Endenicher Allee 60, 53115 Bonn, Germany

## Abstract

Biological evolution of a population is governed by the fitness landscape, which is a map from genotype to fitness. However, a fitness landscape depends on the organisms environment, and evolution in changing environments is still poorly understood. We study a particular model of antibiotic resistance evolution in bacteria where the antibiotic concentration is an environmental parameter and the fitness landscapes incorporate tradeoffs between adaptation to low and high antibiotic concentration. With evolutionary dynamics that follow fitness gradients, the evolution of the system under slowly changing antibiotic concentration resembles the athermal dynamics of disordered physical systems under quasistatic external drives. Specifically, our model can be described as a system with interacting hysteretic elements, and it exhibits effects such as hysteresis loops and memory formation under antibiotic concentration cycling. Using methods familiar from studies in this field, we derive a number of analytical and numerical results. Our approach provides a general framework for studying motifs of evolutionary dynamics in biological systems in a changing environment.

## INTRODUCTION

The concept of the fitness landscape, introduced by Sewall Wright [1], is a useful tool for the visualization of evolutionary processes as populations that are being driven uphill along fitness gradients by natural selection. Mathematically, a fitness landscape is a map from the genotypes of a species to fitness values. In recent decades, it has become possible to empirically determine fitness landscapes for systems comprising multiple mutations, and a wealth of new work has illuminated various aspects of evolutionary dynamics on different kinds of fitness landscapes [2–10]. The theory of fitness landscapes and its application to empirical data has also seen a rise in recent decades [11–16].

A less well-studied topic in this field is the evolution in changing environments. Fitness landscapes are a function of environment, and can change in systematic ways as environmental parameters change. Whereas the fitness landscape provides information about *G* × *G* (gene-gene) interactions, the introduction of the environmental parameter furnishes information about *G* × *G* × *E* (where *E* stands for environment) interactions, i.e., about how the environment modifies the gene-gene interactions [17–20]. A few studies on microbial growth have measured or interpolated fitnesses as a function of environmental parameters [21–23], but systematic theoretical work in this field is still limited.

Understanding and predicting the effect of the environment on fitness landscapes has important practical applications. A pertinent example is the case of antibiotic resistance in bacteria, where it has been shown that the fitness landscape depends strongly on the antibiotic concentration [21, 22]. Uncontrolled variation in antibiotic concentration, both in clinical settings and elsewhere [24, 25], is a cause for the rise in antibiotic resistance, which is a major clinical challenge today. Figure 1 shows an empirical example of the kind of processes we are interested in. The fitnesses of the genotypes in the figure were measured in [21], and based on it, one can predict transitions between genotypes under concentration increase (black/gray arrows) or decrease (red/orange arrows). Notice that this small system already exhibits some interesting properties, such as a hysteresis loop under antibiotic concentration cycling and transient genotypes that are not part of the loop.

**FIG. 1.**
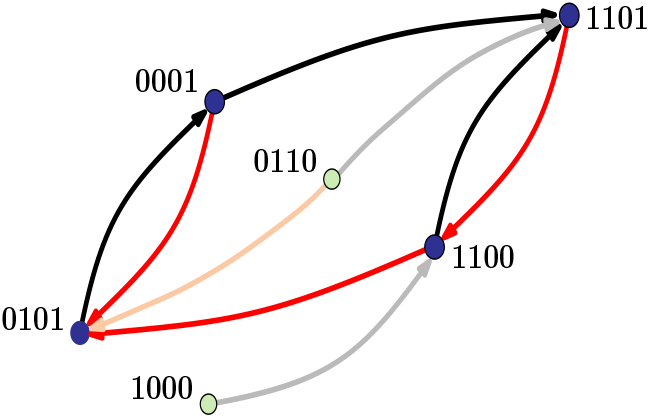
State transition graph of antibiotic resistance evolution. The nodes depict genotypes composed of four mutations in the antibiotic resistance enzyme TEM-50 *β*-lactamase. Genotypes are represented as binary strings where a 1 denotes the presence and 0 the absence of a specific mutation. The growth rates of bacteria expressing these mutant enzymes were reported in [21] for the antibiotic piperacillin at three different concentrations (128 *µ*g/ml, 256 *µ*g/ml and 512 *µ*g/ml). Each node is a local fitness maximum at one of these concentrations. Black and grey arrows connect nodes that would be reached under adaptive evolution when the concentration is increased, and red and orange arrows represent the dynamics under concentration decrease. For example, 0001 is a local maximum at 256 *µ*g ml, but when the concentration is switched to 512 *µ*g/ml, it is no longer a fitness maximum. Evolution through a greedy adaptive walk then leads to the new maximum 1101. The graph displays a hysteresis loop 0101 → 0001 → 1101 → 1100 → 0101. The green nodes cannot be reached under cyclic concentration changes.

Our focus is primarily on one such class of problems, where the environmental dependence of the fitness landscape is governed by a tradeoff between two phenotypes, bacterial growth rate and resistance [23]. While we will mostly use the language of antibiotic resistance evolution in the following, the theory developed is more generally applicable, as will become clear from the mathematical model. Our work uses tools from statistical physics, specifically the physics of disordered systems. Concepts and methods from statistical physics have been used in the theory of evolution for a long time [26, 27]. Precise quantitative analogies with evolutionary phenomena have been found with equilibrium statistical physics [28], the theory of random walks [29], spin glasses [30–32] and many more. Most of these, however, focus on static fitness landscapes. Here we investigate evolution on rugged landscapes that vary with changes in an external parameter. This problem is naturally reminiscent of the physics of driven disordered systems, particularly in the athermal quasistatic (AQS) regime [33], where thermal activation processes are absent or negligible. The primary effect of the external forcing is then to alter the set of stable equilibria or their locations. As a result, under a time-varying external forcing such systems remain in a given equilibrium until it becomes unstable, and a fast relaxation process leads to a new equilibrium. Despite the absence of thermal activation processes, the resulting dynamics can nevertheless be rather complex, exhibiting memory effects [34, 35] as well as dynamic phase transitions, such as the jamming transition in granular materials [36], or the yielding transition in amorphous solids [37].

We find that evolutionary genotypic change has close parallels with systems such as cyclically sheared amorphous solids [38, 39], where a changing environmental parameter is analogous to an external shear, and transitions to new genotypes are similar to localized plastic events inside the solid. As was shown recently [35, 40, 41], the AQS conditions permit a rigorous description of the dynamics of such systems in terms of a directed *state transition graph*. Since the transition graph represents the response of the system to any possible deformation protocol, it provides a bird’s-eye view of the possible dynamics [35, 42, 43]. One of the main goals of this paper is to show that this approach leads not only to new insights into evolution in changing environments, but furnishes new quantitative results through the application of analytical techniques used in the theory of driven disordered systems. We find analogies with certain variations of the Preisach model, which is an elementary model of hysteresis [44] that was introduced in the context of magnetic systems, and has subsequently been generalized by allowing the hysteresis elements, the *hysterons*, to interact with each other in various ways [43, 45–47]. In the following section, we describe our evolutionary model in detail, before moving on to the analogies with disordered systems and their applications.

## MODEL

We define a genotype ***σ*** as a binary string of length *L*, i.e. *σ*_*i*_ ∈ {0, 1}, where *i* = 1, 2, …, *L* denotes the sites where mutations can occur, and *σ*_*i*_ = 1 indicates the presence of a mutation. An equivalent and useful way of thinking about ***σ*** is as a set of mutations drawn from a total of *L* possible mutations. The genotype without mutations, commonly referred to as the *wild type*, is then the empty set, whereas the *all-mutant* is the set with all the *L* mutations. We will use the notation ***σ*** both as a string and as a set, and clarifications on the notation will be provided wherever necessary.

Our focus is on the tradeoff induced landscapes (TIL) model introduced in [23], which is defined through three key properties that are motivated by empirical observations. 1) The fitness of each genotype ***σ*** is a function of an environmental parameter *x* ≥ 0, and the fitness curve has the form

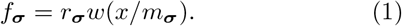

The fitness curve thus has the same shape for different genotypes except for a rescaling of the axes by the genotype-specific parameters *r*_***σ***_ and *m*_***σ***_. We call *r*_***σ***_ the null-fitness and *m*_***σ***_ the resistance of a genotype ***σ***, following terminology used for bacterial *dose-response curves* that represent the population growth rate as a function of drug concentration [23, 48, 49]. We choose units such that for the wild type ***σ*** = **0**, *r*_**0**_ = 1 and *m*_**0**_ = 1, so that *f*_**0**_(*x*) = *w*(*x*). Further, *w*(*x*) is a monotonic decreasing function, reflecting the decreasing fitness of a bacterial cell with increasing drug concentration. 2) Every mutation comes with two parameters *r*_*i*_ and *m*_*i*_, and for any genotype, *r*_***σ***_ = exp[∑_*i*_ *σ*_*i*_ ln *r*_*i*_] and *m*_***σ***_ = exp[∑_*i*_ *σ*_*i*_ ln *m*_*i*_]. Thus, the effects of individual mutations combine in a simple multiplicative manner. 3) We assume that mutations exhibit tradeoff between adaptation to low and high drug concentrations, *i*.*e. r*_*i*_ *<* 1 and *m*_*i*_ *>* 1. This means that every mutation enhances the resistance, but this comes at the cost of reduced null-fitness. The fitness curves of a specific realization of the TIL model with *L* = 2 mutations are shown in Fig 2(a).

**FIG. 2.**
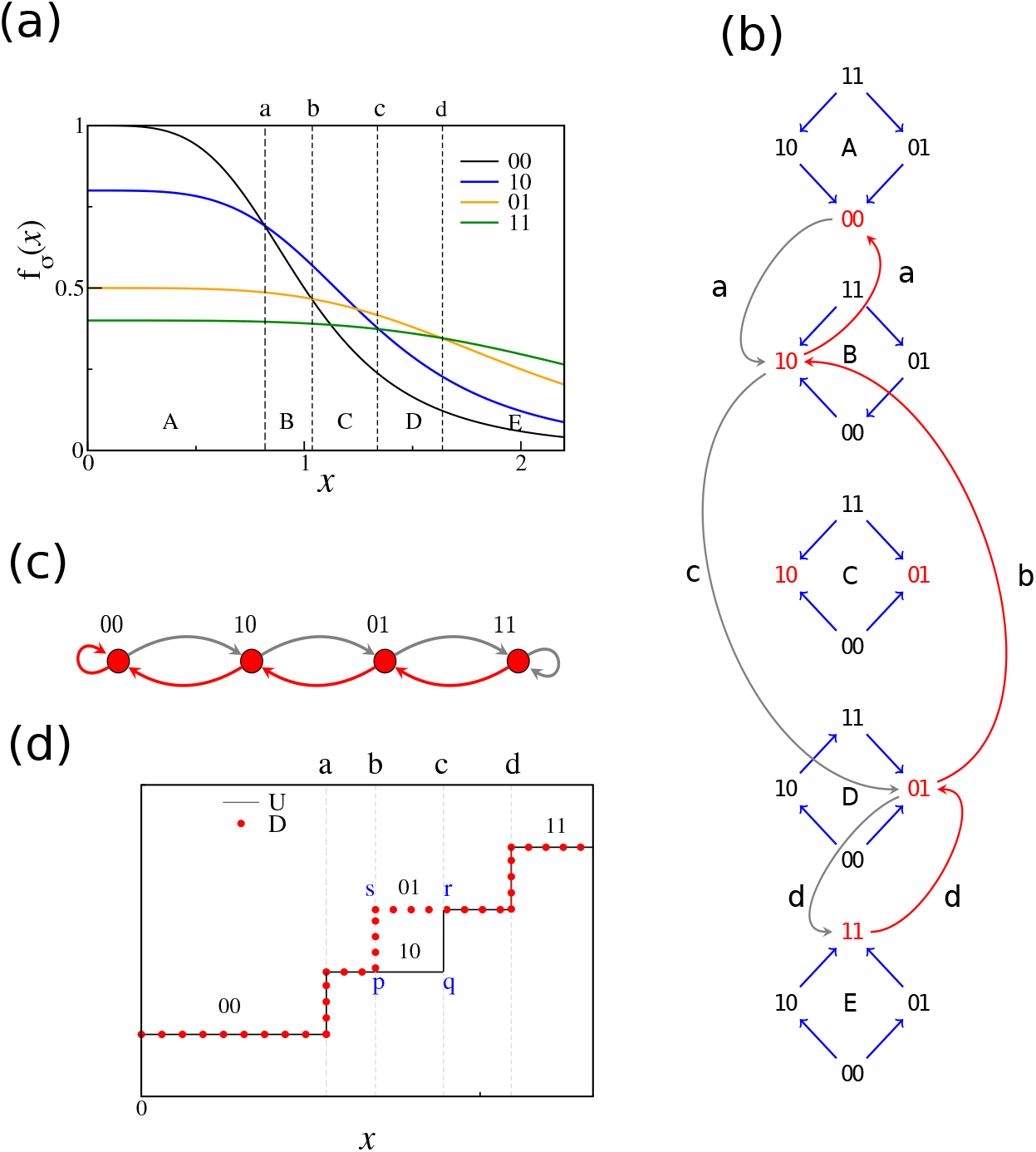
Tradeoff-induced fitness landscapes (TIL) model. (**a**) Fitness curves for four genotypes in a TIL model with two sites (*L* = 2). The parameters are *r*_1_ = 0.8, *m*_1_ = 1.3 and *r*_2_ = 0.25, *m*_2_ = 2.5. The shape of the curve is *w*(*x*) = 1*/*(1 + *x*^2^), which is a Hill function, a form commonly used to model dose-response curves [48, 49]. The figure is divided into five regions A-E, corresponding to different fitness graphs. A new fitness graph occurs when the fitness curves of two mutational neighbors intersect. The *x*-values of the intersection points are marked by the letters a-d. (**b**) Fitness graphs in the regions A-E. Concentration increases in the downward direction. In each fitness graph, the local fitness maxima are marked in red. Evolution in a fitness graph follows the oriented edges until a fitness maximum is reached. The curved grey arrows follow the evolution of the system under quasistatic increase of *x* starting from the stable state 00 at *x* = 0 until the all-mutant 11 is reached; the curved red arrows continue the trajectory as the concentration is quasistatically decreased until 00 is reached again. The points on the *x*-axis at which the transitions occur are stated next to the arrows. Note that not all curved arrows involve changes at a single site. The transition at c alters both sites in two steps, 10 → 11 → 01. (**c**) Transition graph for the two-site system. The nodes are the stable states which, in this simple case, comprise all genotypes. The grey arrows are the U transitions, i.e. the transitions under concentration increase, and the red arrows are the D transitions, i.e. transitions under concentration decrease. The transition graph can be read off from the sequence of fitness graphs in panel (**b**). (**d**) The transition between between states is shown schematically. Each horizontal level is a genotype, and the vertical lines denote transitions. The black lines correspond to genotypes under U transitions (starting from 00 at *x* = 0), while the line traced out by the red dots indicates the genotypes under D transitions (starting from 11 at large *x*). The hysteresis loop *pqrs* is marked in the figure.

The problem of analyzing this model has two components. First, one needs to understand the topography of the fitness landscape, e.g., the set of local fitness maxima and the paths that lead to the maxima, for a fixed *x*. The second part involves questions about evolutionary dynamics between maxima under changing drug concentrations. Specifically, we address scenarios where *x* changes quasistatically, in the sense that, at every value of *x*, evolution reaches a local fitness maximum. The first part has been addressed in detail in [23], and we describe some of the salient features of landscape topography here. The landscape of the TIL model is highly rugged (except at very low and very high *x*), i.e. the number of fitness maxima is asymptotically exponential in *L* [23]. To describe evolutionary dynamics at fixed *x*, it is useful to introduce the notion of a fitness graph. The nodes of the graph are the genotypes, and edges connect mutational *neighbors*, i.e. genotypes that differ by a single mutation. The fitness graph is an acyclic oriented graph, where the edges point towards increasing fitness [16, 50]. Evolution is assumed to proceed through adaptive walks, i.e. the entire population moves along the edges of the fitness graph respecting their orientation [30, 51, 52]. Along the path taken by an adaptive walk the fitness increases monotonically, and such paths are called *(evolutionarily) accessible* [2, 11, 53]. The adaptive walk terminates once a local fitness maximum is reached. In general, there are multiple accessible paths starting from a genotype. We define a *greedy walk* as an adaptive walk at which every step is maximally fitness increasing. It should be clear that the fitness graph is a function of *x*, and a fitness maximum for a certain value of *x* may not be a fitness maximum for another (see Fig. 2(b) for an example). We also make use of the notion of *mutationally directed* (or simply *directed*) paths, which are paths in the fitness graph along which the number of mutations increases or decreases monotonically. The following interesting property about the TIL landscapes at fixed *x* was established in [23]. It is worth discussing, since we will use it to prove certain results in the following sections.

### Directed Path Accessibility: Every (mutationally) directed path ending at a local fitness maximum is accessible

In other words, every local maximum ***σ*** is evolutionarily accessible from the subsets (respectively supersets) of ***σ*** by a sequential gain (resp. loss) of mutations; the mutations may be gained (or lost) in any order. This property has remarkable consequences. For example, the wild type is a subset of every genotype, and therefore can access every fitness maximum through all directed paths. Whenever the wild type is a fitness maximum, it must be the only fitness maximum in the landscape, since it can be accessed from all genotypes. The same two properties also hold for the all-mutant. With this background, we move on to the main focus of this article, which is evolutionary dynamics under slow changes in *x*. It is here that the relation with the Preisach model will become apparent.

## RESULTS

### Stable states

We consider an evolutionary dynamic where the parameter *x* changes quasistatically, i.e. slowly enough that at any *x* the system always reaches a local maximum through an adaptive walk. We call a genotype a *stable state* if it is a local fitness maximum at *some* concentration *x*. As *x* changes, the fitness graph is altered by flipping the direction of one edge every time the fitness curves of two mutational neighbors intersect (see Fig 2(a) and (b)). A stable genotype ***σ*** loses stability once its fitness curve intersects that of a neighbor, and the system transitions to a new stable state by moving along the oriented edges of the new fitness graph. Given a state ***σ***, we define the two disjoint sets *I*^+^[***σ***] = {*i* : *σ*_*i*_ = 1}, *I*^*−*^[***σ***] = {*j* : *σ*_*j*_ = 0}. For *i* ∈ *I*^+^[***σ***], we denote by ***σ***^*−i*^ the configuration obtained form ***σ*** by setting *σ*_*i*_ = 0. Likewise for *j* ∈ *I*^*−*^[***σ***], let ***σ***^+*j*^ denote the configuration obtained from ***σ*** by setting *σ*_*j*_ = 1. Let *x*_*i*_ be the intersection point of the dose-response curves of the wild-type ***σ*** = **0** and the genotype with a single mutation at site *i, i*.*e*. **0**^+*i*^. Hence, *x*_*i*_ is the solution of the equation

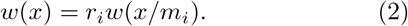

By a suitable choice of the function *w*(*x*) this solution can be guaranteed to be unique (see [23] and *SI*). It then follows that for ***σ*** and *i* ∈ *I*^*−*^[***σ***], the fitness curves of ***σ*** and ***σ***^+*i*^ intersect at *m*_***σ***_ *x*_*i*_. Likewise, for *j* ∈*I*^+^[***σ***] the fitness curves of ***σ*** and ***σ***^*−j*^ intersect at 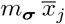, where we have defined 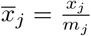. We now see that a necessary and sufficient condition for a genotype ***σ*** to be a stable state is that

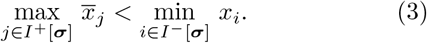

If this holds, let the index, *ℓ* (*u*) correspond to the site where the maximum (minimum) on the left (right) hand side of the inequality is attained. The stability range of ***σ*** is then 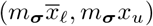.

At this point, we introduce a fruitful analogy with the standard Preisach model, which is comprised of a set of non-interacting two-level systems referred to as *hysterons* [44, 54]. The mutation variable *σ*_*i*_ ∈ {0, 1} is analogous to the *i*-th hysteron, and its states 0 and 1 to the up and down states of the hysteron. The parameter *x* plays the role of an external magnetic field that drives the system. In the Preisach model, each hysteron has an upper (lower) threshold at which it transitions to the up (down) states as the magnetic field reaches the threshold from the lower (upper) direction. We define the *Preisach analogue* of the TIL model as composed of *L* hysterons, where the upper and lower thresholds of the *i*-th hysteron are *x*_*i*_ and 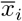 respectively. Note that since *m*_*i*_ *>* 1, we have 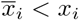. There is hysteresis since the transition from the down to the up state occurs at a higher value of the field than the transition from up to down. Referring to Eq. 3 we arrive at our first key result:

#### The stable states of the TIL model and its Preisach analogue are identical

However, as we will show next, there are significant differences in the dynamical properties of the two models.

### Dynamics and the transition graph

As mentioned before, the dynamics under quasistatically changing *x* can be described by transitions among stable states. We call a transition under concentration increase a U-transition, and that under concentration decrease a D-transition. Then the dynamics can be described by a *transition graph* (see Fig 2(c) for an example) where the nodes are the stable states, and each node has outgoing U and D edges. While the TIL model and its Preisach analogue share the same set of stable states, the dynamical properties are in general different. To illustrate this, in Fig 3 we show a particular realization of the TIL model with *L* = 5 along with its Preisach analogue. In the Preisach model, each transition comprises of a single switching of the least stable hysteron, which leads to a new stable state. In the TIL model, a change at a single site need not lead to a stable state. For example, the U-transition 15 → 27 in the TIL model in Fig 3 involves changes at the third and fifth sites.

**FIG. 3.**
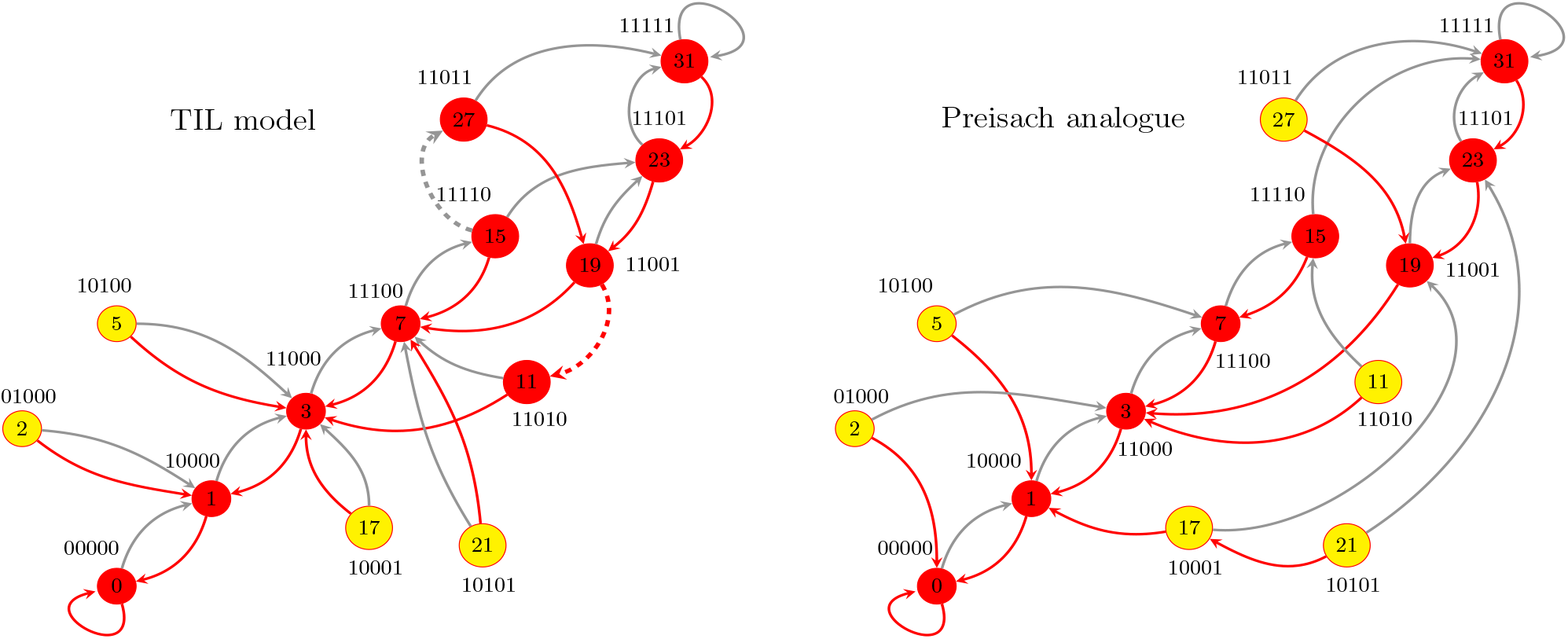
Transition graph of a realization of the TIL model (left) and its Preisach analogue (right) with *L* = 5 sites. The null-fitness and resistance parameters (*r*_*i*_, *m*_*i*_) of the 5 mutations were drawn randomly from a distribution specified in the *SI*. The symbolic ordering sequence of this realization is given in Eq. 4. Each genotype is assigned an integer label, placed within the nodes, by interpreting the genotype string as a binary code where the leftmost digit is the least significant. The red arrows are U-transitions and grey arrows are D transitions. The yellow nodes are the genotypes that cannot be reached starting from the wild type. When multiple outgoing arrows are present from a state, the solid ones correspond to greedy walks, i.e maximal fitness increase at every intermediate step, whereas the dashed lines represent fitness-increasing walks but with one or more steps that are not maximally fitness-increasing.

To understand why, first notice that under concentration changes in the U direction, the first change in a stable state ***σ*** (which must satisfy Eq. 3) is a flip 0 → 1 at the site *u* which has the smallest *x*_*i*_ among sites with *σ*_*i*_ = 0. This flip occurs at *x* = *m*_***σ***_*x*_*u*_. The new state ***σ***^+*u*^ also satisfies Eq. 3 (since 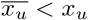) and is therefore a stable state. However, in order for ***σ***^+*u*^ to be a local fitness maximum at *x* we must require that its lower stability threshold is less or equal to *x*. This threshold is 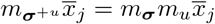, where *j ∈ I* ^+^[***σ***^+*u*^] and 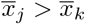 for all *k ≠ j* with *k* ∈ *I*^+^[***σ***^+*u*^]. Therefore the new state is a fitness maximum if and only if 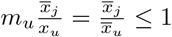, in which case the dynamics terminates at ***σ***^+*u*^. When the condition is violated, additional *secondary mutations* occur until a fitness maximum is reached. While many detailed properties of the secondary mutations depend on system parameters, certain features are general and in particular do not depend on the choice of *w*(*x*). In the following we will mention some of these.

In the Preisach model, exactly *L* U-transitions are required to get from the wild type to the all mutant, and exactly *L* D-transitions to go from the all-mutant to the wild type, as is seen in Fig 3. In the TIL model, due to the existence of secondary mutations, these numbers are generally different. The TIL model in Fig 3 requires 6 U-transitions to go from the wild type to the all-mutant, and 6 D-transitions in the reverse direction, even though *L* = 5. One important consequence of the secondary mutations is that *the number of mutations does not always increase monotonically under U or decrease monotonically under D*. The first secondary mutation is always of a complementary kind to the initial mutation, where the changes 0 → 1 and 1 → 0 are defined to be of complementary kind to each other (see *SI* for a proof). Further mutations may also continue to be complementary to the initial mutation, leading to a (temporary) decrease in the number of mutations under U or an increase under D, as shown in a typical trajectory for *L* = 20 mutations in Fig 5(a). This seems counterintuitive, but arises from the state-dependent pre-factor *m*_***σ***_ in the stability thresholds of stable states.

Moreover, when secondary mutations are present, the state ***σ′*** to which a transition occurs from a state ***σ*** need not be unique, due to the possible presence of multiple adaptive paths. In Fig 3, the state 15 can transition either to the state 27 or the state 23 under concentration increase. It can also be shown, using the property of direct path accessibility, that secondary mutations cannot cause a transition to a subset or superset of ***σ*** (see *SI*), i.e. both the initial and the final state must contain at least one mutation not contained in the other. Another related consequence of the secondary mutations is that ***σ*** may transition to the same state ***σ′*** under U and D-transitions. For example, the state 5 in the TIL graph in Fig 3 transitions to the state 3 under both U and D-transitions. This also appears counterintuitive from a biological standpoint, but it can occur when the stability range of ***σ*** is contained in that of ***σ***′.

To understand the transition graph of the TIL model in a more systematic way, we adopt a strategy that has been fruitful for the Preisach model [54]. We construct a symbolic sequence *p* that specifies the total order among all the elements of 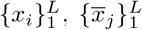. First, without loss of generality, we order our indices *i* such that *x*_1_ *< x*_2_ *< … < x*_*L*_. Next, it is useful to define the permutation *ρ* of (1, 2, …, *L*) that orders the 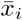 among themselves from largest to smallest, so that 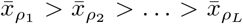. Since 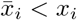 for each *i*, what remains is the specification of the ordering relation between 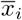 and *x*_*j*_ for *j≠i*. Given the sets 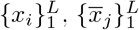 and the ordering prescribed by *ρ*, we can describe the total ordering in terms of a symbolic sequence *p* of elements 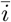 and *i* by making the correspondence 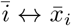 and *i ↔x*_*i*_, so that the sequence specifies the increasing order of *x*_*i*_ and 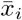. Since 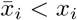 and the permutation *ρ* have to be respected, the sequence has to be such that the following hold: for each *i*, 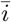 is to the left of *i*; the subsequence of sites without overbars is 1, 2, …, *L*; the subsequence of sites with overbars is *ρ*_*L*_, *ρ*_*L−*1_, …, *ρ*_1_. As an example, consider the TIL Model in Fig 3, which has *L* = 5 and *ρ* = (43521), and the ordering

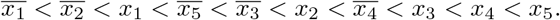

The corresponding symbolic ordering sequence *p* is then

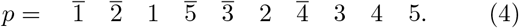

Because of the Preisach-TIL correspondence, whether a genotype/Preisach state is stable or not, and what the least stable sites of a stable state are, can be read off from *p*, since the condition in Eq. 3 is easy to check by inspecting *p*. In the case of the Preisach model this implies that *p* completely determines the transition graph [54]. While this is not the case for the TIL model, the sequence *p* nevertheless contains considerable information about the TIL transition graph. In particular, this representation provides a precise condition for the existence of secondary mutations (see *SI*):

#### Secondary mutations are absent from all transitions in the TIL model if and only if the ordering sequence is of the form

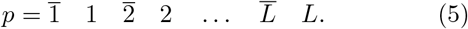

In the absence of secondary mutations, the transition graph of the TIL model becomes identical to that of its Preisach analogue. The transition graph in this case has a simple chain structure (see Fig 4), and the number of stable states is *L*+1, which is the lowest possible in a TIL (or Preisach) model. Note that despite identical transition graphs in this case, some dynamical differences are still present. Each Preisach element is hysteretic, and therefore forward and reverse transitions between two states do not occur at the same concentration; in the TIL model satisfying Eq. 5, however, they occur at the same concentration, namely the one at which the dose-response curves of the two genotypes intersect.

**FIG. 4.**
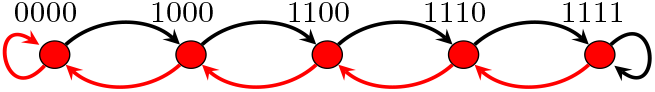
TIL transition graph with *L* = 4 and no secondary mutations. The ordering sequence is of the form given in Eq. 5. The transition graph is unique for this ordering sequence, and is identical to the graph for the Preisach analogue.

### Hysteresis, reversibility and memory

The reversibility of evolution under a reversal of environmental conditions is an important question in evolutionary biology [55–57] In the specific case of antibiotic resistance evolution, to what extent resistance is reversed in a drug-free environment is a question of considerable clinical importance [58–60] One should note that different notions of reversion are used here. One common definition refers to a sudden (rather than slow) change in environment to a new state, followed by a switch back to the original state [58, 61, 62]. In the context of our model, we study reversion under quasistatic environmental changes, which would appear to be most conducive for approximately reversible behavior. The phenomenon is naturally linked to the notion of hysteresis under a slow and continuous change of an external field, and indeed the Preisach model was first proposed as a simplified, tractable model of hysteresis [44, 54].

It is clear that the TIL model, in general, also exhibits hysteresis and irreversibility. The highest degree of reversibility is exhibited by systems with chain-like transition graphs, such as in Fig 2(c) or Fig 4, where each transition ***σ*** *→* ***σ***′ is accompanied by the transition ***σ***′ *→* ***σ*** in the reverse direction, and there are no states with multiple outgoing edges in either direction. This means that under a reversal of the direction of concentration change, the same genotypes occur in reversed sequence. However, the transitions ***σ*** *→* ***σ***′ and ***σ*** ′ *→* ***σ*** need not occur at the same concentration. For example, in Fig 2(d), the transition 10 *→* 01 occurs at the point *x* = *c* during concentration increase, but 01 *→* 10 occurs at *x* = *b* during concentration decrease. On the other hand, for systems of the type shown in Fig 4, the forward and reverse transitions occur at the same concentration. However, such perfect reversibility is not typical of TIL models. In general, TIL graphs have forward transitions with no corresponding reverse transitions, such as the D-transition 11 *→* 3 or the U-transition 15 *→* 27 in the TIL graph in Fig 3. Based on the observation of the systems described in Fig 2 and 4, we need to distinguish between two kinds of hysteresis loops. We say that two states ***σ*** and ***σ***′ form a *concentration loop* if one can go from ***σ*** to ***σ***′ under quasistatic concentration increase and from ***σ***′ to ***σ*** under concentration decrease, and there is some range of *x* over which the forward and reverse trajectories do not share the same genotype. The system in Fig 2 exhibits a concentration loop, as shown by the rectangle with corners marked by the points *p, q, r*, and *s* in Fig 2(d). We say that ***σ*** and ***σ***′ form a *graph loop* (***σ***,***σ***′) if one can go from ***σ*** to ***σ***′ under U-transitions and from ***σ*** to ***σ***′ under D-transitions, and if there is at least one genotype contained in either the forward or reversed sequence of states that is not contained in the other. A graph loop between two states implies a concentration loop, but a concentration loop does not imply a graph loop. For example, the case shown in Fig 2 has a concentration loop, but does not have a graph loop, as can be readily seen in Fig 2(c). An example of a graph loop in the TIL model is the one formed by the U transitions from 7 to 23 and the D transitions from 23 to 7. A necessary condition for the existence of graph loops in the TIL model is the presence of secondary mutations; for otherwise, every transition must be among mutational neighbors, and such that the upper stability threshold of one coincides with the lower stability threshold of the other, causing the transitions to be reversible.

Hysteresis is also linked to the notion of memory [34, 35]. A genotype encountered along a trajectory not only contains information about the concentration, but also about the history of environmental change. At the simplest level, it may contain information about whether one is on the U or D boundary of a loop. For example, in the region between *x* = *b* and *x* = *c* in Fig 2(d), the state 10 indicates that the concentration has been increasing, while 01 indicates that it was decreasing. But there is more information available than this, in general. The subloops seen in the TIL graph in Fig 3 contain (partial) information about extreme values of *x* reached in previous rounds of concentration cycling. Thus, if the dynamics started with the wild type at *x* = 0 and reached the genotype 11 at some point, one infers that the last transition happened by a lowering of the concentration to below the stability threshold of 19; but we also see from the transition graph that on some previous upward path the concentration must have exceeded the upper stability threshold of 15, followed by some sequence of transitions that brought it to the lower stability threshold of 19 for the first time since this happened.

In this context, it is important to mention the phenomenon of *return point memory* (RPM) [40, 63] possessed by certain systems, which in our setting of adaptive evolution implies that genotypes at which the direction of the concentration change has been reversed can be returned to with a subsequent reversal and hence remembered. The RPM property is universally present in the Preisach model [34, 40, 54]. In the context of state transition graphs, one talks about the loop-RPM property, which ensures that the system cannot escape any loop between two states ***σ*** and ***σ***′ without passing through one of these states (see [40, 54] for a detailed exposition). The RPM property implies the loop-RPM property, and is therefore possessed by the Preisach model and can be checked for the Preisach graph in Fig 3.

Return-point memory is a mechanism by which a memory of local extremes of the driving parameter can be retained. For example, in the TIL model and under greedy dynamics, starting with the wild type at *x* = 0, and increasing the concentration until state 15 is reached, any decrease of concentration followed by a subsequent increase will eventually lead again to state 15. However, if the concentration continues to increase, so that state 23 is reached, then a concentration decrease to 19 followed by an increase will not lead to 15 anymore. Thus the memory of 15 as the genotype at a local extreme event has been erased and replaced by 23. While we have found many realizations of the TIL model that possess the loop-RPM property under greedy transitions, it is not universally present. Additionally, the existence of alternative fitness increasing transitions, such as the ones shown in Fig 3 by the dashed lines, can cause a loss of this kind of memory. To see this, consider the loop formed by the greedy U transitions leading from state 7 to 23 and the greedy D transitions going from 23 to state 7. On the downward trajectory from 23, it is possible to escape the loop without going first through 7 by making the transition 19 *→* 11.

### Statistical properties of the main hysteresis loop

While the properties of the TIL model described so far depend only on the ordering sequences *ρ* and *p* encoded by the transition graph, for a more detailed study of the concentration-dependent evolutionary dynamics the fitness values of the model need to be explicitly assigned. In the standard Preisach model, one usually considers the thresholds to be independent random variables. Similarly for the TIL model, we assume that *r*_*i*_ and *m*_*i*_ follow a joint probability distribution with density *P* (*r, m*) and the ordered pairs (*r*_*i*_, *m*_*i*_) for *i* = 1, 2, … *L* are independently and identically distributed. Their joint probability density is then given by 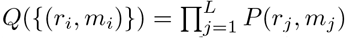.

Since *x*_*i*_ and 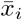 are functions of *r*_*i*_ and *m*_*i*_ only, the pairs 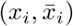 for *i* = 1, 2, … *L* are independently and identically distributed as well. Let the (marginal) cumulative distribution function (CDF) of *x*_*i*_ be *F*_*x*_(*x*_*i*_) and that of 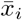 be 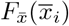, and let 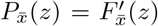 denote the probability density function of 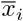. The probability that a genotype is a fitness maximum is the probability that Eq. 3 holds. The calculation of this probability is facilitated by the fact that *I*^+^[***σ***] and *I*^*−*^[***σ***] are disjoint sets. One can show that in the limit of large *L* the average number of stable states ⟨*N*_*ss*_⟩ is given by

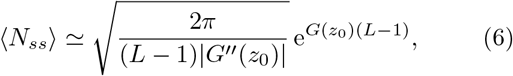

where the average is taken with respect to 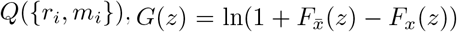), and the global maximum of this function is at *z*_0_. The mean number of states is asymptotically exponential in *L*, showing the highly rugged nature of these landscapes (see *SI* for the derivation of Eq. 6).

An important dynamical question is understanding the evolutionary sequence of genotypes as the concentration is cycled between very low and very high values. Investigating this is facilitated by considering only greedy U and D transitions. We numerically generate trajectories starting from the wild type at *x* = 0 and increasing *x* quasistatically until the all-mutant is reached, and then decreasing *x* quasistatically until the wild type is reached again. We call this trajectory the *main hysteresis loop*, and the upward and downward parts of it the U-and D-boundary respectively. The mean number of mutations in the genotype as a function of *x* on both the U and D boundaries are shown in Fig 5(a), which clearly shows hysteresis. The inset shows a comparison with the Preisach model. Note that in the TIL model the range of concentrations over which a genotype ***σ*** is a local maximum has an overall scale factor *m*_***σ***_. Therefore, in order to facilitate comparison of the data for the TIL model and its Preisach analogue, we have rescaled the concentration axis for the former by ⟨*m*(*x*)⟩, the average scale factor *m*_***σ***_ of the states ***σ*** that are stable at concentration *x*. From the inset of 5(a) we see that for the Preisach model the curve for the number of mutations on the U-boundary is seen to be lower than the corresponding curve for the D-boundary. This effect can be understood qualitatively as follows. The intersection points along the U-boundary are governed by the distribution of *x*_*i*_ and those along the D-boundary by the distribution of 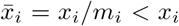. As a result, the *i*-th mutation is acquired at a larger *x* along the U-boundary compared to where it is lost on the D boundary. More generally, for any randomly chosen pair of mutations *i* and *j, x*_*j*_ tends to be higher than 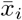 since all the *m*_*i*_’s are larger than 1. The consequence is that the intersections along the U-boundary tend to occur at larger values of *x* compared to the D-boundary, making the the curve for the number of mutations along the U-boundary lower. Essentially the same effect is visible for the TIL model when the *x* values are rescaled by the value of ⟨*m*(*x*)⟩ on the boundaries. When the rescaling is not done, the U-boundary becomes higher, as seen in the main Fig 5(a). The clue to understanding this comes from Fig 5(b), which shows that the average resistance level ⟨*m*(*x*)⟩ at given *x* is lower for the U-boundary. Since the intersection points have the pre-factor *m*_*σ*_ in the TIL model, this effect tends to make the intersection points along the U-boundary occur at lower values of *x*. For our system, this effect is apparently strong enough to shift the curve of the number of mutations along the U boundary above that of the D-boundary.

**FIG. 5.**
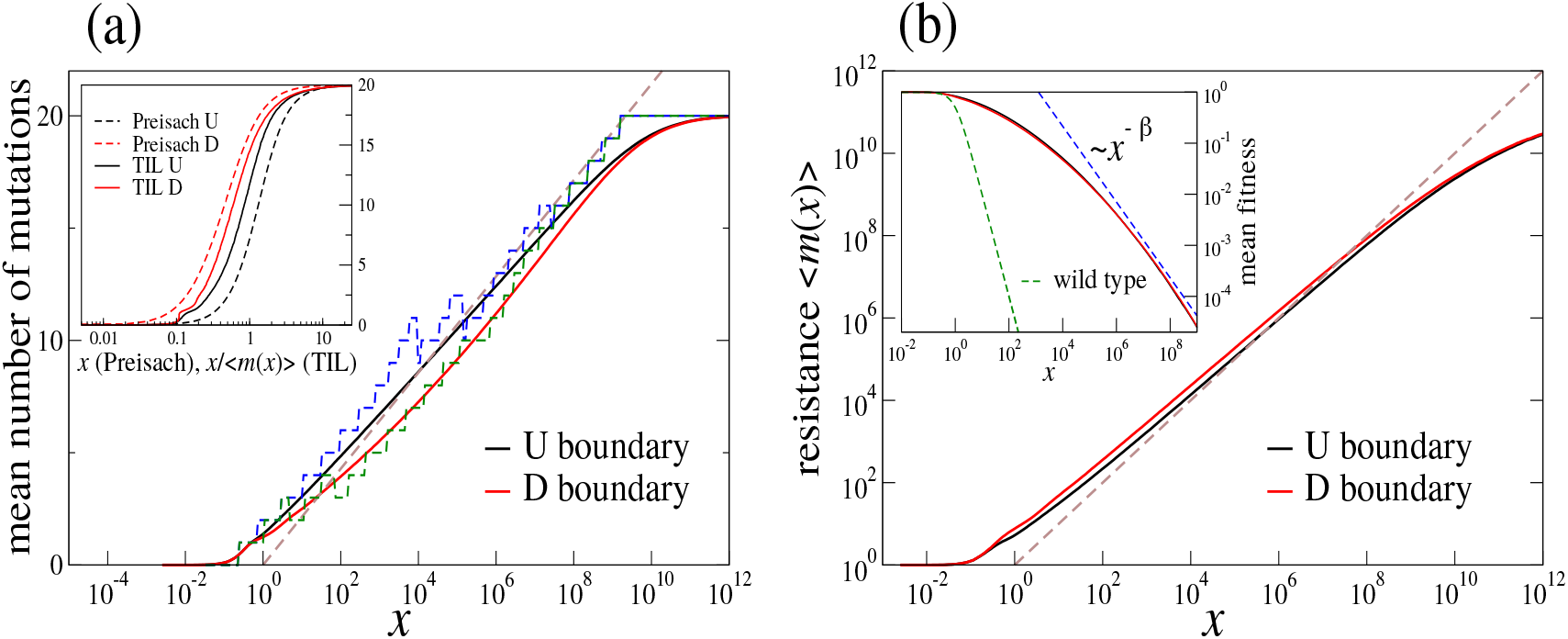
Average properties of greedy evolutionary trajectories along the main hysteresis loop for *L* = 20. The dose-response curve is *w*(*x*) = 1*/*(1 + *x*^2^). The mutation parameters (*r*_*i*_, *m*_*i*_) are drawn randomly from a joint probability density described in the *SI*. A total of 10^4^ realizations were used to calculate averages. (**a**) Mean number of mutations in the genotypes along the U and D boundaries, which describe the behavior under increasing (U) and decreasing (D) concentrations. The dashed blue and green lines show a typical sample trajectory. The dashed brown line is the curve 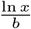, where *b* = ln ⟨*m*⟩. The inset shows rescaled values of the mutation number in comparison to the Preisach analogue (dashed lines). The rescaling is done by the mean resistance level ⟨*m*(*x*)⟩. (**b**) Mean resistance level ⟨*m*(*x*)⟩ of the genotypes along the boundaries. The brown dashed curve is *x*. The inset shows the fitness of the genotypes along the boundaries. The dashed green line is the fitness of the wild type *w*(*x*), and the dashed blue line is the power law *x*^*−β*^, where 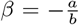 and *a* = *−⟨*ln *r⟩*. Angular brackets denote averages with respect to the distribution *Q*(*{*(*r*_*i*_, *m*_*i*_)*}*) (see main text for further details).

Generally, the changes of resistance level and mutation number have a complex mutual dependence, and these can vary between systems depending on the dose response curve and the parameter distribution. However, certain asymptotic features that hold generically for stable states can be computed to leading order. For example, the mean number of mutations in a state that is stable at *x* scales asymptotically as 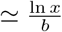, where the parameter *b* = ⟨ln *m*⟩, and *m* is the resistance of an individual mutation. This is indicated by the brown dashed line in Fig 5(a). At the same level of approximation, the mean level of resistance satisfies the relation ln ⟨*m*(*x*)⟩ ≃ ln *x*, which is shown as a dashed brown line in Fig 5(b). The inset of Fig 5(b) displays the fitness as a function of *x*, which is seen to decline at a much lower rate than that of the wild type, as a consequence of the increasing level of resistance. Detailed derivations of these results are given in the *SI*.

### Secondary mutations and path irreversibility

Figure 6 shows simulation results for the mean number of secondary mutations as a function of the number of background mutations (i.e., the number of mutations in the genotype from which the transition originates) along the main hysteresis loop. The curves appear to become largely independent of *L* at large *L*. Moreover, for large *L*, the number of secondary mutations depends weakly on the number of background mutations (unless the latter is close to 0 or *L*). Overall the number of secondary mutations is seen to be well below 1. Therefore, even for large *L*, the greedy adaptive walks at fixed *x* are seen to be short (provided, of course, one starts from genotypes on the main loop that are very close to their stability range).

**FIG. 6.**
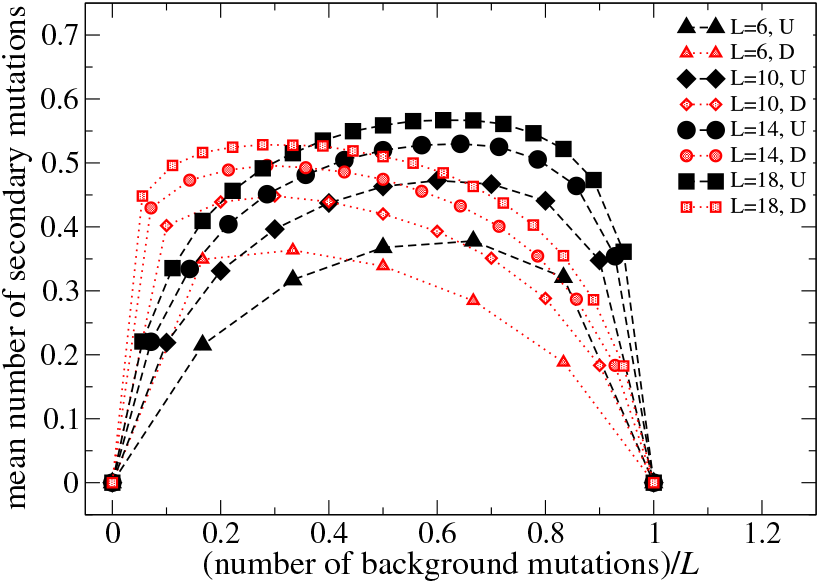
Mean number of secondary mutations in an evolutionary transition as a function of the number of mutations in the originating background genotype. Simulations were conducted along the main hysteresis loop using greedy dynamics, and the mutation parameters were generated randomly as described in the text and the *SI*. We used 10^5^-10^6^ realizations. Black symbols correspond to the U boundary and red to the D boundary.

As was shown above, secondary mutations are also the source of genotypic irreversibility in the TIL model. We now describe a measure of irreversibility, adapted from a distance measure for evolutionary paths introduced in [64]. Let ***σ*** be the genotype at *x* on the U boundary of the main loop. Then *d*_*U*_ (*x*) is defined as the minimum of the Hamming distance between ***σ*** and the genotypes on the D boundary (for any *x*). The quantity *d*_*D*_(*x*) can be defined in an analogous way. These quantities are plotted in Fig 7. The distance measures vanish at very low and high concentrations, which is expected since the wild type and all-mutant are on both the U and D boundaries of the loop. The maximum value is reached close to the concentration at which the estimated mean number of mutations is *L/*2, as shown by the vertical dotted lines. The maximum value of the distance reached for *L* = 18 is about 1.4. The level of reversibility is thus high, consistent with the low number of secondary mutations, and the relatively narrow hysteresis loops seen in Fig 5.

**FIG. 7.**
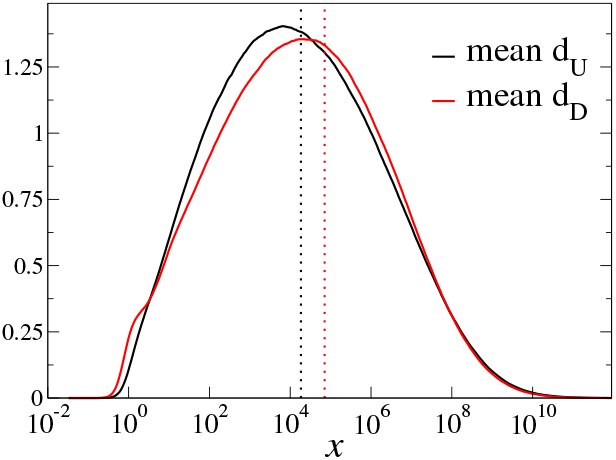
Mean Hamming distance between genotypes on the U and D boundary as a function of concentration for L=18. A total of 10^5^ realizations were used to calculate averages. The distance measure *d*_*U*_ (*x*) (*d*_*D*_ (*x*)) is the minimal Hamming distance between the current genotype on the U (D) boundary and any point on the D (U) boundary. A low value corresponds to high reversibility. The black (red) dotted line is the concentration at which the mean number of mutations is *L/*2 on the U (D) boundary, obtained from Fig 5(a).

## SUMMARY

We have investigated a class of models of bacterial evolution under changing drug concentrations, and shown that their dynamics are closely related to that of driven disordered systems, exhibiting dynamical phenomena such as hysteresis and memory formation. We have applied analytical tools developed within the context of driven disordered systems in order to characterize the evolutionary dynamics. The transitions between genotypes in a homogeneous population are found to exhibit a number of generic properties which we have described in detail. Adaptive walks are found to be short, i.e they involve a small number of mutations, when a slow change in the drug concentration renders a local fitness maximum unstable. Moreover, the systems generically exhibit hysteresis loops, i.e a lack of reversibility when the direction of concentration change is reversed at certain points in the trajectory. However, the degree of reversibility is found to be rather high, which is related to the quasistatic nature of change in concentration studied here. We have also obtained asymptotic scaling approximations to the number of mutations, and to the fitness and resistance level of genotypes, which are expected to be relatively robust to modifications in the driving protocol or the precise assumptions of the model. Conceptually, our analysis highlights the distinction between reversibility on the level of genotypes and on the level of phenotypes such as resistance or fitness.

We have shown that partial information about the changing environment is stored in evolving genotypes, and can be recovered from state transition graphs. Although our analysis has focused on the relatively simple TIL model, we should emphasize that the memory effects described here are generic features of disordered systems. The analogies presented in this paper are therefore indicative of a much greater scope for the study of information about the past history of a population that is embedded in its genotype as it is evolves under changing environmental conditions.

The approach used in this paper can be easily extended to systems that go beyond the specific assumptions of the model investigated here. Of particular interest is the extension to situations where the environment, and hence the fitness landscape, changes periodically in discontinuous, discrete steps. An important realization of this scenario is the cycling of different antibiotics in the treatment of bacterial infections, where fitness landscape theory can be used to devise treatment plans that are optimized to avoid or slow down resistance evolution [65, 66]. This problem will be pursued in future work.

## I. ACKNOWLEDGEMENTS

SGD and JK acknowledge support by the Deutsche Forschungsgemeinschaft (DFG, German Research Foundation) within SFB 1310 *Predictability in evolution*. MM was supported by the DFG under project No. 398962893, project No. 211504053-SFB 1060, and under Germanys Excellence Strategy-GZ 2047/1, project No. 390685813.

## SUPPLEMENTARY INFORMATION

This file includes details of the mathematical models and computations used in the main text.

## MATHEMATICAL DETAILS

We first derive a series of general results for the TIL model which are independent of the shape of the dose-response function *w*(*x*) as long as it satisfies the following properties:

(W1) *w*(*x*) is a continuous, strictly decreasing function for *x* ≥ 0.

(W2) For all pairs of permissible values (*r, m*) such that *r* < 1 and *m* > 1, the curves *w*(*x*) and *rw*(*x/m*) intersect at precisely one point.

As noted in [23] these conditions are satisfied for a large class of dose-response functions considered in the literature, including the exponential and half-Gaussian functions. When considering statistical results, we will specialize to the *n* = 2 Hill-type dose-response function *w*(*x*) = 1*/*(1 + *x*^2^). In this case, in order for (W2) to be satisfied, the permissible pairs (*r, m*) have to satisfy also *m*^2^*r* > 1, as is readily checked.

As given in the main text, the stability condition of a state is

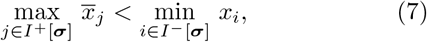

In the following, we shall assume that we are given a stable genotype ***σ*** such that

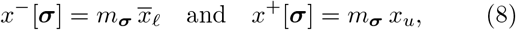

hold with *x*^−^[***σ***] < *x*^+^[***σ***], implying that the stability condition Eq. 7 is satisfied, and the sites *ℓ* and *u* are those at which the maximum, respectively minimum of the terms on the left and right hand side of the inequality are attained. We say that sites *ℓ* and *u* are the least stable sites in the sense that the first mutation will occur there under concentration decreases or increases, respectively. The following two statements are readily shown. Given any pair of sites *ℓ* and *u* such that 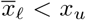, there exists a symbolic ordering sequence *p* such that Eq. 8 holds and thus *ℓ* and *u* are the least stable sites for some stable state ***σ***. Conversely, given a symbolic ordering sequence *p*, using Eq. 7, the set of all stable genotypes ***σ*** can be inferred from it, and hence the possible pairs of least stable sites (*ℓ, u*) associated with these. Whenever we assume that Eq. 8 holds, this will either imply that we are given a specific order sequence *p* and that with respect to *p* the state ***σ*** is stable with (*ℓ, u*) being the pair of least-stable sites, or alternatively, we are given (*ℓ, u*) and consider the set of order sequences *p* and stable states ***σ*** compatible with this choice of least stable sites. The particular point of view will be clear from the context.

### Secondary mutation must be complementary

We first derive some simple inequalities for the TIL model. The following is readily shown from the assumptions (W1) and (W2) made above for *w*(*x*):

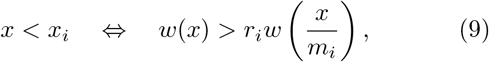

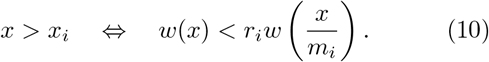

Now, let ***σ*** be a stable state, such that Eq. 8 holds. Then, we can show from the previous results that for all *i* ∈ *I*^−^[***σ***] \ {*u*},

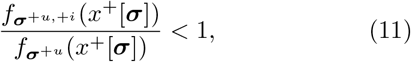

and for all *j* ∈ *I*^+^[***σ***] \ {*ℓ*},

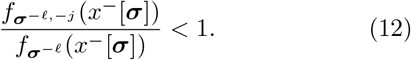

The last two inequalities together assert that first secondary mutations which are in the same direction as the primary mutation, are fitness-decreasing. Thus either the first mutation leads to a local fitness maximum, and hence there will be no further mutations, or the first secondary mutation must be complementary to the original mutation.

### Locations of complementary secondary mutations

Let ***σ*** be a stable state such that Eq. 8 holds. Then for *i* ∈ *I*^+^[***σ***],

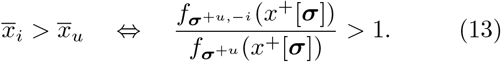

Therefore, subsequent to an initial mutation under concentration increase at site *u*, fitness increasing complementary mutation sites are those sites *i* ∈ *I*^+^[***σ***] for which the symbol 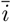 is located to the right of *ū* in the order sequence *p*. Note in particular, that the initial mutation site *u* itself cannot be also the site for a subsequent secondary mutation, as this would have implied that ***σ*** has a higher fitness than ***σ***^+*u*^ at the triggering concentration.

Likewise, for *j* ∈ *I*^−^[***σ***],

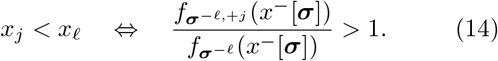

Any secondary mutation following an initial mutation under concentration decrease at site *ℓ*, must be a site *j* ∈ *I*^−^[***σ***] located to the left of *ℓ* in the symbolic order sequence *p*. The statements Eq. 13 and Eq. 14 are proven by repeated application of Eq. 9, Eq. 10, and the properties of ordering sequences *p* that are compatible with the assumption Eq. 8.

### Secondary mutations cannot cause transitions to a subset or superset

Assume the contrary. Then, according to the property of Directed Path Accessibility, a path must exist where the first secondary mutation is in the same direction as the original mutation. But this is not possible according to the previous result.

### Conditions for the absence of secondary mutations

Consider a realization of the TIL model with *L* sites and let ***σ*** be a stable state satisfying Eq. 8. Assume that we are given a symbolic ordering sequence *p* compatible with Eq. 8. For any site *u* = 1, 2, … *L*, we will be interested in the interval of elements of *p* that is bounded to the left by *ū* and to the right by *u*. Denote by ℐ_*u*_ the set of sites *j* that appear in this interval without overbars. Likewise, let 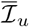 be the set of sites that appear in this interval with overbars. Our definition is such that neither of the two sets of sites ℐ_*u*_ and 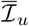 contain *u*.

Consider now transitions out of ***σ*** under concentration increases. By assumption, under a concentration increase to (a value slightly above) *x*^+^[***σ***], the site *u* will mutate first, *σ*_*u*_ = 0 → 1, leading to ***σ***^+*u*^, and as a result, the upper limit of the stability range of ***σ***^+*u*^ increases to *x*^+^[***σ***^+*u*^] > *x*^+^[***σ***]. In order to assert the stability of ***σ***^+*u*^ at the concentration *x*^+^[***σ***] which triggered the mutation at *u*, we must require that

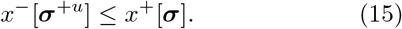

If this condition is not satisfied, then *x*^*±*^[***σ***^+*u*^] > *x*^+^[***σ***] and at least one secondary mutation occurs. We thus need to find conditions under which Eq. 15 holds.

Now in terms of the ordering sequence *p*, the site 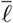 must be located to the left of *u*, as must be the site *ū*. Moreover, since *ℓ* and *u* have to be distinct, 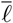 is either to the left or right of *ū*. In the former case we have 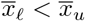, and hence

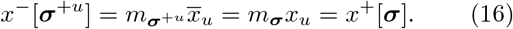

Since Eq. 15 is satisfied, genotype ***σ***^+*u*^ is a local fitness maximum at this concentration and there will therefore be no secondary mutations. Suppose next that 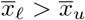. In this case

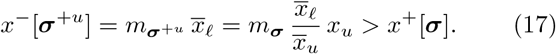

Therefore there will be at least one complementary secondary mutation at some site *i* ∈ *I*^+^[***σ***]. Condition Eq. 13 asserts that in order for such a mutation to be fitness increasing, *i* must be such that 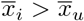. Using Eq. 7, it is easily shown that the stability condition of ***σ***, as given by Eq. 8, implies that 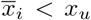, so that the secondary mutation site must be contained in the set 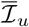. Note in particular that the site, *e* itself satisfies these conditions and hence is a possible candidate for the first secondary mutation. Combining all of the above results, under increasing concentration, a secondary mutation will occur if and only if the set 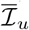 is non-empty, and the state ***σ*** is such that, for some 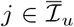 we have *σ*_*j*_ = 1. Conversely, a secondary mutation will *not* occur if and only if one of the following two conditions holds:

(U1) The set 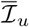 is empty.

(U2) The set 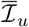 is non-empty, and the state ***σ*** is such that, for each 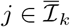 we have *σ*_*j*_ = 0.

In a similar manner, one can show that under decreasing concentration a secondary mutation will not occur, if and only if one of the following two conditions holds:

(D1) The set ℐ_*ℓ*_ is empty.

(D2) The set ℐ_*ℓ*_ is non-empty, and the state ***σ*** is such that, for each *j* ∈ ℐ_*ℓ*_ we have *σ*_*j*_ = 1.

Observe now that in order for secondary mutations to be absent from all transitions in a TIL model, the sets ℐ_*k*_ and 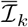 have to be empty for each *u* = 1, 2, …, *L*, since otherwise there will exist stable states for which conditions (U2) or (D2) can be made not to hold. As is easily shown, the only ordering sequence for which both of these sets are empty for each *u* is the sequence

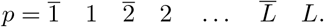

### Asymptotic number of stable states

Consider a genotype ***σ*** with *n* mutations, i.e. ∑ _*i*_ ***σ***_*i*_ = *n*. The number of such genotypes is 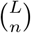 and such a ***σ*** is a stable state if Eq. 7 holds. Since *x*_*i*_ and 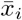 are independent for distinct sites, the probability density that the left hand side of Eq. 7 is less than *z* and the right hand side is greater than *z* is 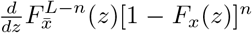. Then the mean number of stable states is

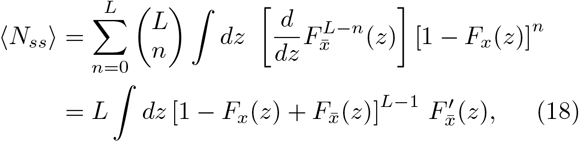

from which the result in the main text follows using a saddle point approximation for *L* large.

### Probability density function used in the numerics

We assume that the dose-response curve is of Hill-type with *n* = 2. In order to satisfy the requirement (W2) for the dose-response function, the parameters (*r*_*j*_, *m*_*j*_) must be chosen such that 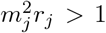 for each *j* = 1, 2, …, *L*. We further assume that the pairs (*r*_*j*_, *m*_*j*_) are independently and identically distributed, so that their joint density is given by 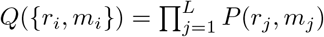. We write *P* (*r*_*j*_, *m*_*j*_) = *P*_1_(*r*_*j*_)*P*_2_(*m*_*j*_|*r*_*j*_). We chose

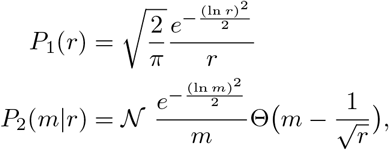

where Θ(·) is the Heaviside step function, and 𝒩 is the appropriate normalization constant.

### Asymptotic approximation for number of mutations

The fitness *f* of a genotype ***σ*** can be expressed as

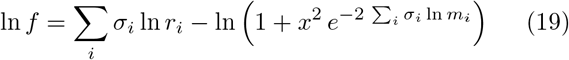

The number of mutations in the genotype is *n* = Σ _*i*_ ***σ***_*i*_. A simple heuristic that produces good approximations for the mean of various quantities at large *L* is a s follows: we consider the fitness of a genotype to be a function of *x* and *n* only, and replace the parameters associated with the mutations by suitable averages. Thus, we write Eq. 19 as

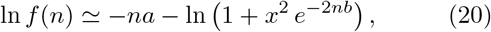

where *a* = − ⟨ln *r*⟩ and *b* = ⟨ln *m*⟩. For any given *x*, one can now maximize Eq. 20 with respect to *n*, yielding an approximation to the mean mutation number at *x* for stable maxima. Taking the derivative of the above with respect to *n* and setting it to zero produces the equation:

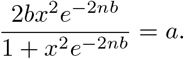

The solution to this is

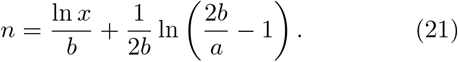

For large *x* and therefore large *n*, the leading order is

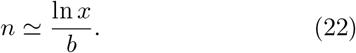

This estimate works well when *L* is large and 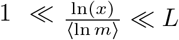.

## Notes

### Competing Interest Statement

The authors have declared no competing interest.

## References

[1] S. Wright, in Proc. Sixth. Int. Cong. Genet., Vol. 1 (1932) pp. 356–366.

[2] D. M. Weinreich, N. F. Delaney, M. A. DePristo, and D. L. Hartl, Science 312, 111 (2006).

[3] E. R. Lozovsky, T. Chookajorn, K. M. Brown, M. Imwong, P. J. Shaw, S. Kamchonwongpaisan, D. E. Neafsey, D. M. Weinreich, and D. L. Hartl, Proc. Natl. Acad. Sci. USA 106, 12025 (2009).

[4] M. F. Schenk, I. G. Szendro, M. L. M. Salverda, J. Krug, and J. A. G. M. de Visser, Molecular biology and evolution 30, 1779 (2013).

[5] J. A. G. M. de Visser and J. Krug, Nature Reviews Genetics 15, 480 (2014).

[6] A. C. Palmer, E. Toprak, M. Baym, S. Kim, A. Veres, S. Bershtein, and R. Kishony, Nature communications 6, 1 (2015).

[7] C. Bank, S. Matuszewski, R. T. Hietpas, and J. D. Jensen, Proc. Natl. Acad. Sci. USA 113, 14085 (2016).

[8] J. Domingo, G. Diss, and B. Lehner, Nature 558, 117 (2018).

[9] I. Fragata, A. Blanckaert, M. A. D. Louro, D. A. Liberles, and C. Bank, Trends in Ecology & Evolution 34, 69 (2019).

[10] V. O. Pokusaeva, D. R. Usmanova, E. V. Putintseva, L. Espinar, K. S. Sarkisyan, A. S. Mishin, N. S. Bogatyreva, D. N. Ivankov, A. V. Akopyan, S. Y. Avvakumov, I. S. Povolotskaya, G. J. Filion, L. B. Carey, and F. A. Kondrashov, PLoS Genetics e1008079, 15 (2019).

[11] J. Franke, A. Klözer, J. A. G. M. de Visser, and J. Krug, PLoS Computational Biology 7, e1002134 (2011).

[12] I. G. Szendro, M. F. Schenk, J. Franke, J. Krug, and J. A. G. M. de Visser, Journal of Statistical Mechanics: Theory and Experiment 2013, P01005 (2013).

[13] J. Neidhart, I. G. Szendro, and J. Krug, Genetics 198, 699 (2014).

[14] L. Ferretti, B. Schmiegelt, D. Weinreich, A. Yamauchi, Y. Kobayashi, F. Tajima, and G. Achaz, Journal of theoretical biology 396, 132 (2016).

[15] F. Blanquart and T. Bataillon, Genetics 203, 847 (2016).

[16] K. Crona, A. Gavryushkin, D. Greene, and N. Beeren-winkel, eLife 6, e28629 (2017).

[17] A. Murugan, K. Husain, M. J. Rust, C. Hepler, J. Bass, J. M. Pietsch, P. S. Swain, S. G. Jena, J. E. Toettcher, A. K. Chakraborty, et al., Physical Biology 18, 041502 (2021).

[18] M. G. J. de Vos, S. E. Schoustra, and J. A. G. M. de Visser, Europhysics Letters 122, 58002 (2018).

[19] F. A. Gorter, M. G. M. Aarts, B. J. Zwaan, and J. A. G. M. de Visser, Genetics 208, 307 (2018).

[20] D. W. Anderson, F. Baier, G. Yang, and N. Tokuriki, Nature Communications 12, 3867 (2021).

[21] P. M. Mira, J. C. Meza, A. Nandipati, and M. Barlow, Molecular biology and evolution 32, 2707 (2015).

[22] C. B. Ogbunugafor, C. S. Wylie, I. Diakite, D. M. Weinreich, and D. L. Hartl, PLoS computational biology 12, e1004710 (2016).

[23] S. G. Das, S. O. Direito, B. Waclaw, R. J. Allen, and J. Krug, Elife 9, e55155 (2020).

[24] D. W. Kolpin, M. Skopec, M. T. Meyer, E. T. Furlong, and S. D. Zaugg, Science of the Total Environment 328, 119 (2004).

[25] D. I. Andersson and D. Hughes, Nature Reviews Microbiology 12, 465 (2014).

[26] R. A. Fisher, The genetical theory of natural selection (Dover, 1958).

[27] E. V. Koonin, The logic of chance: the nature and origin of biological evolution (FT press, 2011).

[28] G. Sella and A. E. Hirsh, Proceedings of the National Academy of Sciences 102, 9541 (2005).

[29] M. Manhart and A. V. Morozov, in First-passage phenomena and their applications (World Scientific, 2014) pp. 416–446.

[30] S. Hwang, B. Schmiegelt, L. Ferretti, and J. Krug, Journal of Statistical Physics 172, 226 (2018).

[31] D. L. Stein, Spin glasses and biology (World Scientific, 1992).

[32] S. Franz, L. Peliti, and M. Sellitto, J. Phys. A: Math. Gen. 26, L1195 (1993).

[33] C. E. Maloney and A. Lemaître, Phys. Rev. E 74, 016118 (2006).

[34] N. C. Keim, J. D. Paulsen, Z. Zeravcic, S. Sastry, and S. R. Nagel, Rev. Mod. Phys. 91, 035002 (2019).

[35] M. Mungan, S. Sastry, K. Dahmen, and I. Regev, Phys. Rev. Lett. 123, 178002 (2019).

[36] R. P. Behringer and B. Chakraborty, Reports on Progress in Physics 82, 012601 (2018).

[37] D. Bonn, M. M. Denn, L. Berthier, T. Divoux, and S. Manneville, Rev. Mod. Phys. 89, 035005 (2017).

[38] I. Regev, T. Lookman, and C. Reichhardt, Phys. Rev. E 88, 062401 (2013).

[39] D. Fiocco, G. Foffi, and S. Sastry, Phys. Rev. E 88, 020301 (2013).

[40] M. Mungan and M. M. Terzi, Ann. Henri Poincaré 20, 2819 (2019).

[41] M. Mungan and T. A. Witten, Phys. Rev. E 99, 052132 (2019).

[42] I. Regev, I. Attia, K. Dahmen, S. Sastry, and M. Mungan, Phys. Rev. E 103, 062614 (2021).

[43] N. C. Keim and J. D. Paulsen, arXiv preprint 2101.01240 (2021).

[44] F. Preisach, Zeitschrift für Physik 94, 277 (1935).

[45] O. Hovorka and G. Friedman, Journal of magnetism and magnetic materials 290, 449 (2005).

[46] C. W. Lindeman and S. R. Nagel, arXiv preprint 2101.01632 (2021).

[47] M. van Hecke, arXiv preprint 2107.06556 (2021).

[48] R. R. Regoes, C. Wiuff, R. M. Zappala, K. N. Garner, F. Baquero, and B. R. Levin, Antimicrobial agents and chemotherapy 48, 3670 (2004).

[49] G. Chevereau, M. Dravecká, T. Batur, A. Guvenek, D. H. Ayhan, E. Toprak, and T. Bollenbach, PLOS Biology 13, e1002299 (2015).

[50] K. Crona, D. Greene, and M. Barlow, Journal of Theoretical Biology 318, 1 (2013).

[51] S. Kauffman and S. Levin, Journal of theoretical Biology 128, 11 (1987).

[52] A. Agarwala and D. S. Fisher, Theor. Pop. Biol. 130, 13 (2019).

[53] D. M. Weinreich, R. A. Watson, and L. Chao, Evolution 59, 1165 (2005).

[54] M. M. Terzi and M. Mungan, Phys. Rev. E 102, 012122 (2020).

[55] H. Teotónio and M. R. Rose, Evolution 55, 653 (2001).

[56] J. T. Bridgham, E. A. Ortlund, and J. W. Thornton, Nature 461, 515 (2009).

[57] M. Kaltenbach, C. J. Jackson, E. C. Campbell, F. Hollfelder, and N. Tokuriki, eLife 4, e06492 (2015).

[58] D. I. Andersson and D. Hughes, Nature Reviews Microbiology 8, 260 (2010).

[59] R. C. Allen, J. Engelstädter, S. Bonhoeffer, B. A. McDonald, and A. R. Hall, Proceedings of the Royal Society B: Biological Sciences 284, 20171619 (2017).

[60] P. Durão, R. Balbontín, and I. Gordo, Trends in microbiology 26, 677 (2018).

[61] A. Dunai, R. Spohn, Z. Farkas, V. Lázár, Á. Györkei, G. Apjok, G. Boross, B. Szappanos, G. Grézal, A. Faragó, et al., Elife 8, e47088 (2019).

[62] P. S. Zur Wiesch, J. Engelstädter, and S. Bonhoeffer, Antimicrobial agents and chemotherapy 54, 2085 (2010).

[63] J. P. Sethna, K. Dahmen, S. Kartha, J. A. Krumhansl, B. W. Roberts, and J. D. Shore, Phys. Rev. Lett. 70, 3347 (1993).

[64] A. E. Lobkovsky and E. V. Koonin, Frontiers in Genetics 3, 246 (2012).

[65] P. M. Mira, K. Crona, D. Greene, J. C. Meza, B. Sturmfels, and M. Barlow, PloS ONE 10, e0122283 (2015).

[66] D. Nichol, P. Jeavons, A. G. Fletcher, R. A. Bonomo, P. K. Maini, J. L. Paul, R. A. Gatenby, A. R. Anderson, and J. G. Scott, PloS Comp. Biol. 11, e1004493 (2015).

